# Establishment and validation of a High-throughput Micro-Neutralization assay for Respiratory Syncytial Virus (subtypes A and B)

**DOI:** 10.1101/2023.03.20.533425

**Authors:** Carolina Bonifazi, Claudia Maria Trombetta, Irene Barneschi, Simona Latanza, Sara Leopoldi, Linda Benincasa, Margherita Leonardi, Claudia Semplici, Pietro Piu, Serena Marchi, Emanuele Montomoli, Alessandro Manenti

**Author notes:** Corresponding author: Serena Marchi, Department of Molecular and Developmental Medicine, University of Siena, via Aldo Moro 2, 53100 Siena, Italy. Equal contribution (Carolina Bonifazi, Claudia Maria Trombetta). (CB), (CMT), (SM), (EM). (IB), (SL), (SLE), (CS), (PP), (AM). (LB), (ML). **Statements**. Funding: This research received no external funding. Ethics approval: Not applicable. Patient consent: Not applicable. Permission to reproduce material from other sources: Not applicable. Clinical trial registration: Not applicable.

## Abstract

The validation of a bioanalytical method allows us to determine its validity for a designated purpose and to guarantee the reliability of its analytical results. The virus neutralization assay has proved to be suitable for the detection and quantization of specific serum neutralizing antibodies against respiratory syncytial virus subtypes A and B. Respiratory syncytial virus is a negative-sense RNA virus and is responsible for the majority of acute lower respiratory tract infections in infants and older adults worldwide. Owing to its widespread infection, the WHO considers it a target for the development of preventive vaccines. Despite the high impact of its infections, however, no vaccine candidate is currently available.

The aim of this paper is to provide a detailed validation process for the micro-neutralization assay and to demonstrate that this method can effectively support the efficacy assessment of candidate vaccines and the definition of correlates of protection.

## INTRODUCTION

Respiratory syncytial virus (RSV) is the most important respiratory tract pathogen of early childhood (1). The virus is estimated to cause 33 million cases of disease and 66,000 to 199,000 deaths every year in children under 5 years of age worldwide (2). Most of the children affected by RSV are aged 2 years or less and repeat infections can occur throughout life (3). The virus can also be dangerous for older adults, patients with chronic heart or lung disease and those with a weakened immune system (4).

RSV belongs to the *Paramyxoviridae* family, *Pneumovirinae* subfamily, which also includes human metapneumovirus, however it is the only member of the genus *Pneumovirus* able to infect humans (1, 5). It is an enveloped, non-segmented single negative-strand RNA virus; the genome contains 10 genes encoding 11 proteins. Among these, the three surface viral proteins are the attachment (G) glycoprotein, the fusion (F) glycoprotein and the small hydrophobic (SH) protein. The G and F proteins are the main surface glycoproteins and are crucial for the infectivity and pathogenesis of the virus. The G protein enables the virus to attach to the respiratory epithelial cells of the host, while the F protein causes the fusion of viral and host cell membranes and, in a later stage, the fusion of infected cells, thereby inducing the characteristic syncytia, from which the name of the virus is derived. The F protein exists in two different conformations: pre-fusion (pre-F) and post-fusion (post-F). The pre-F conformation is a metastable form that is necessary in order to initiate viral entry. The pre-F conformation is rich in neutralization-specific sites (Ø and V), which, by contrast, are not present in the post-F conformation (6). The G protein also contains virus neutralization epitopes, which, along with F, are usually the designated targets of therapeutic drugs and vaccines (1, 5, 7, 8). On the basis of its antigenic and genetic variability, RSV has been classified into two subtypes, A and B. Both subtypes can circulate simultaneously during annual epidemics, but RSV-A prevails over RSV-B in most years (5, 8). The F protein is well conserved in RSV-A and B, their amino acid sequences being 90% identical or higher (5, 9).

Before the COVID-19 pandemic, RSV infections showed a seasonal trend in temperate regions, with peaks during winter months (10). The public health and social measures taken to mitigate the impact of the pandemic led to a reduction in RSV or even its disappearance. On the other hand, however, they contributed to increasing the number of naïve subjects, including that of naïve older children. RSV-neutralizing antibodies (nAb) were seen to be reduced in childbearing and breastfeeding women, increasing the number of children susceptible to RSV (7, 11-13). Interestingly, studies (11, 14-20) have reported low levels of RSV circulation following the outbreak of SARS-CoV-2, but also an out-of-season outbreak following the relaxation of public health measures, due to the accumulation of susceptible subjects. By contrast, a parallel increase in RSV infection in older adults was not observed, suggesting that behavioral changes implemented during the pandemic may have reduced RSV infection rates in this age-group (21).

Currently, there is no vaccine to prevent RSV, though many candidates are in clinical development. It seems that the failure, in terms of protection and, above all, of safety, of the first formalin-inactivated RSV vaccine in 1960 has overshadowed the study and development of RSV vaccines for almost 40 years (22). The only available preventive measure is prophylaxis with Palivizumab, a humanized monoclonal anti-RSV antibody that is recommended for use in highest-risk infants (23). This aspect highlights the importance of nAb in protecting against severe disease, as is supported by evidence that high levels of nAb correlate significantly with protection against RSV challenge in adult volunteers (24, 25).

The virus neutralization (VN) assay is a serological technique that can detect the presence of functional nAb capable of inhibiting virus replication (26). This feature makes the VN assay particularly suitable for evaluating the immunogenicity of RSV vaccines. However, the lack of standardized assays is a major drawback (8). The introduction of an international reference standard -the WHO 1^st^ IS for Antiserum to RSV has improved assay harmonization across formats, allowing readings to be converted to international units (IU) per milliliter and facilitating comparison of the performance of candidate vaccines even when different assay formats are used (8, 27).

The present study aimed to document the results obtained during the validation of the VN assay against RSV-A and B in accordance with the International Council for Harmonization (ICH) guidelines (28). The objective was to demonstrate the suitability of this assay for the detection and quantization of serum-nAb, with a view to testing clinical samples in trials evaluating the immunogenicity of new vaccines.

## MATERIALS AND METHODS

### Vero cell culture

Vero cells (American Type Culture Collection [ATCC] #CCL-81) were cultured in Minimum Essential Medium with Earle’s salts (EMEM) (Euroclone, Pero, Italy) supplemented with 10% v/v Fetal Bovine Serum (FBS) (Euroclone, Pero, Italy), 2 mM L-glutamine, 100 units/mL penicillin, 100 µg/mL streptomycin (P/S) (Gibco, Life Technologies) with the addition of 1% v/v MEM Non-Essential Amino Acids Solution 100X (Gibco, Life Technologies).

### Cell and Virus propagation

HEp-2 cells (American Type Culture Collection [ATCC] #CCL-23) were cultured in high-glucose Dulbecco’s Modified Eagle’s Medium (DMEM) (Euroclone, Pero, Milan) supplemented with 10% v/v FBS (Euroclone, Pero, Milan), 2 mM L-Glutamine, 100 units/mL penicillin, and 100 µg/mL Streptomycin (P/S) (Gibco, Life Technologies). Cells were maintained at 37°C, in a humified 5% CO_2_ environment, and passaged every 3-4 days.

Human RSV-B WV/14617/85 and human RSV-A were purchased from the American Type Culture Collection (ATCC Number: VR-1400™ and ATCC Number: 1540™, respectively). Viral propagation was performed in 175 cm^2^ tissue-culture flasks pre-seeded with 50 mL of HEp-2 cells (1.0 ×10^6^ cells/mL) diluted in DMEM 10% FBS. After 18-24 hours’ incubation at 37°C in 5% CO_2_, flasks were washed twice with sterile Dulbecco’s phosphate buffered saline w/o Calcium w/o Magnesium (DPBS) (Euroclone, Pero, Italy) and then inoculated with the RSV-A and B viruses at a multiplicity of infection (MOI) of 0.003. The sub-confluent cell monolayer was incubated with the virus for 2 hours at 37°C in 5% CO_2_; flasks were then filled with 50 mL of DMEM 2% FBS and incubated at 37°C in 5% CO_2_. Cells were monitored daily until 20-30% of cytopathic effect (CPE) and 60-70% of syncytial formation were observed. The supernatant was removed from the flasks and collected, the cells were scraped, pooled with the supernatant and centrifuged at 469 g for 5 minutes at 4°C to separate the cells from the viral solution. The supernatant was collected, the cell pellet was resuspended in 1 mL of the supernatant, and three freeze-thaw cycles were performed by placing the vial with the cell pellet on ice; the vial was then placed in a 37°C water bath and vortexed (all steps were carried out for 30 seconds) (29). The supernatant previously collected was added to the cell pellet, mixed gently, aliquoted and stored at -80°C in the presence of sucrose (30).

### Serum Samples

The experimental set-up for the validation of the Micro-neutralization (MN) ELISA-based test used the commercially available human and animal serum samples reported in **Table S1**. We used the antiserum to Respiratory Syncytial Virus WHO 1st International Standard (NIBSC) as a homologous control sample for Specificity experiments, and sheep antisera to Influenza Anti-A/Michigan/45/2015 (H1N1) (NIBSC), Anti-A/Hong Kong/4801/2014 (H3N2) (NIBSC) and Anti-B/Brisbane/60/2008 (B/Victoria) (NIBSC) as heterologous samples. A pool of normal human sera purchased from Discovery Life Sciences was used as a positive sample for Dilutional Linearity experiments. Finally, depleted human serum lacking IgA/IgG/IgM (Sigma-Aldrich) was used as a negative control.

### Live-Virus Micro-Neutralization assay

The setup and validation experiments are described below. The MN tests were performed in 96-well, flat-bottomed, tissue culture, microtiter plates (31). Homologous and heterologous serum samples (human serum samples had been previously heat inactivated at 56°C for 30 min) were placed in the first well of the microtiter plates at a final concentration of 1:20, and serial two-fold dilutions were performed. All dilutions were performed in MEM medium (Euroclone) with 2% FBS, and the final volume was 50 μL per well. 100 TCID_50_ (50% tissue culture infectious dose) of virus in 50 μL of MEM with 2% FBS was then added to each well, and the mixture was incubated for 1 h at 37°C in a humidified CO_2_ incubator. At the end of the incubation time, 3.0 × 10^4^ Vero cells in 100 μL of MEM with 2% FBS were added to each well, and the plates were incubated at 37°C in a humidified CO_2_ incubator. After 3 days of incubation, the plates were emptied and hand-washed twice with DPBS. Fixation was performed by adding 100 μL of a cold 80% vol/vol solution of Acetone (Sigma-Merck, St. Louis, MO, USA) in DPBS and incubating the plates for 10 min at room temperature (RT). After the incubation period, the plates were emptied and air dried. The ELISA test was performed on the same day as fixation. Plates were washed 3 times by means of an automatic plate washer (each well was washed with 300 μL/well of wash buffer prepared with DPBS + 0.3% of Tween 20 (Sigma-Merck)) and 100 μL/well of the primary antibody solution was added. The primary antibody, Anti-RSV Antibody, clones 133-1H, 131-2G, and 130-12H (Sigma-Merck) was diluted in a 1:10,000 ratio by using the antibody diluent (a 5% non-fat dried milk solution in wash buffer). Plates were incubated for 1 hour at RT. Subsequently, the plates were washed three times, as previously, and 100 μL of the secondary antibody solution - Goat Anti-Mouse IgG (H+L)-HRP Conjugate (Bio-Rad, Hercules, CA, USA) diluted in a 1:2,000 ratio in the antibody diluent - was then added and the plates were incubated for 1 hour in the dark at RT. Next, the plates were automatically washed for the third time (6 rinses with 300 µl of wash buffer per well) and 100 μL of the substrate solution was added to each well. The substrate solution had previously been prepared by adding one o-Phenylenediamine dihydrochloride tablet (OPD, Sigma-Merck) to 20 mL of a citrate buffer. The plates were incubated for 10 minutes at RT in the dark. The reaction was stopped by adding 100 μL of stop solution (2.8% vol/vol Sulfuric Acid (Sigma-merck) in distilled water) and the plates were read at an optical density of 490 nm (OD490) by means of a SpectraMax ELISA plate (Medical Device) reader.

### Validation Parameters and Statistical Analysis

Each sample was tested in 4 different analytical sessions run by two operators over two days with three replicate measurements per session. For each of the three replicate measurements, a geometric mean titer (GMT) was calculated. The validation testing design is shown in **Figure S1**.

#### Dilutional Linearity

Linearity was assessed by testing the sample in a 2-fold dilution scheme in which at least one dilution had a titer below the lower limit of quantitation of the assay, starting from a dilution of 1:20. Thus, as the sample in the first well was diluted 1:20, the starting dilution in the first well was 1:20, 1:40, 1:80, 1:160, 1:320, 1:640, 1:1280, 1:2560, 1:5120, 1:10240, 1:20480 and 1:40960. The above-mentioned sample dilutions were tested in one repetition per plate, in three different plates by two different operators on day 1 and day 2.

The parameter was evaluated by examining the relationship between the base-2 logarithm of the GMT (observed titers) and the base-2 logarithm of the serum dilutions across the factorial design. The coefficient of determination (R^2^), y-intercept and slope of the regression line were calculated and reported.

#### Relative Accuracy

The accuracy of the test was evaluated by using the reportable (RP) values obtained during the evaluation of Linearity. The accuracy can be tested by using either a conventional true value or an accepted reference value (28). The GMT of the expected values was calculated from the GMT of the results obtained from the neat sample, and by dividing this value by the corresponding factor of the 2-fold serial dilution.

Relative accuracy was evaluated by calculating the percentage of recovery on the GMT of the RP values and the expected (true) titer and applying the formula: 100*(GMT observed / GMT expected).

#### Precision

Precision was assessed by using the results yielded by the linearity tests. Three aspects of precision were considered: repeatability, intermediate precision and format variability.

##### Precision – Repeatability

Intra-run variability, or repeatability, is the variation expected across replicates under the same operating conditions over a short period of time (28).

##### Precision – Intermediate Precision

Intermediate precision is determined from the total variance component. It indicates the variations and random events that can occur within laboratories, such as days, environmental conditions, operators and equipment.

##### Precision – Format Variability

Format variability represents the variation expected across GMT results yielded by multiple replicates in routine testing. For its calculation, we considered two independent runs consisting of one replicate.

#### Limit of Quantitation and Range

The lower (LLOQ) and upper (ULOQ) Limits of Quantitation were determined as the lower and upper 95% CI of the observed GMT of the lowest and highest sample concentrations at which the assay yielded linear, accurate and precise results.

#### Specificity

Specificity was assessed by testing anti-homologous and anti-heterologous serum samples. The positive sample for the homologous strain had to show a 4-fold difference from the heterologous strains tested.

#### Robustness

The robustness of an analytical procedure is its capacity to remain unaffected by small but deliberate variations during measurements and gives an indication of its reliability.

Two critical conditions were evaluated: cell suspension concentrations and virus-serum mixture incubation time.

## RESULTS

### Set-up of the RSV propagation

The virus propagation setup for both RSV subtypes used two different cell lines, HEp-2 and Vero cells (32). Initially, both lines were widely used to study RSV infection, proving permissive to RSV and sensitive to different RSV subtypes, as shown in **Figure 1** (33, 34).

**Figure 1:**
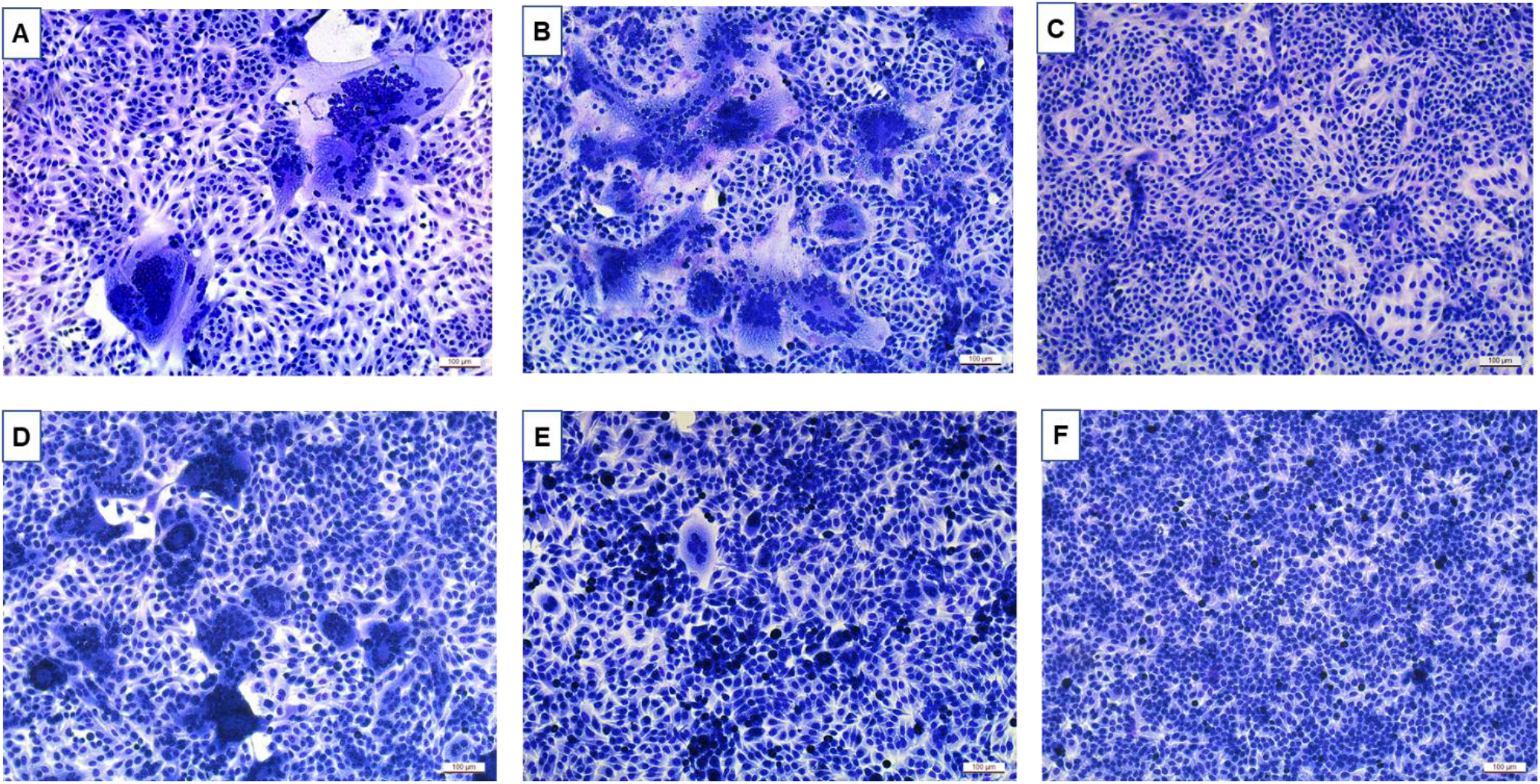
Different pictures of syncytia caused by RSV A infection in Vero cell line (A) and HEp-2 cell line (D), by RSV B infection in Vero cell line (B) and HEp-2 cell line (E) and control cells in Vero cell line (C) and HEp-2 cell line (F).

However, as documented in various studies, the Vero cell-grown virus can infect human airway epithelial cell cultures 600-fold less efficiently than the HEp-2 cell-grown virus (35).

In addition, we observed a fluctuating loss of infectivity by both RSV subtypes, which can be explained by particle instability and aggregation caused by freeze-thawing and handling (29). It has been demonstrated that many sugars, of which sucrose seems to have a more impactful effect, are able to preserve the viability of viruses during freeze-thaw cycles and to avoid RSV aggregation (36). The best MN titers with the highest stability were obtained with HEp-2 cell-grown virus and the optimal sucrose concentration was 3%.

### Setup and validation of MN assay

Before the validation experiments, a series of preliminary analyses was performed in order to select the optimal conditions. To this end, different types of cell lines were used - Vero and HEp-2 - at different concentrations (150,000 and 300,000 cell/mL); both cell suspension and cell adhesion were then tested. The use of different cell media was also investigated by using both the MEM w/ Earle’s Salts w/o L-Glutamine medium (MEM) and DMEM. The most suitable MN incubation time was then investigated on days 2, 3, 4 and 5. Finally, a cross test was performed in order to find the best primary and secondary antibody concentrations. The optimal combination was: Vero at a concentration of 300,000 cell/mL, cell suspension with MEM medium and 3-day incubation. The best concentration of the primary antibody was 1:10,000, while that of the secondary HRP conjugated anti-mouse IgG was 1:2,000.

### Linearity

Dilution linearity is assessed in order to demonstrate that a high concentration of the sample of interest can be diluted to a concentration within the working range and still give a reliable result (37).

The MN assay is considered to have acceptable linearity if, for each RSV subtype, the 95% confidence interval of the slope is between 0.7 and 1.3. In our study, R^2^ was close to 1 for both RSV-A and B, whereas the absolute values of the slope were 1.014 and 1.001, respectively. Results are reported in **Figure 2**.

**Figure 2:**
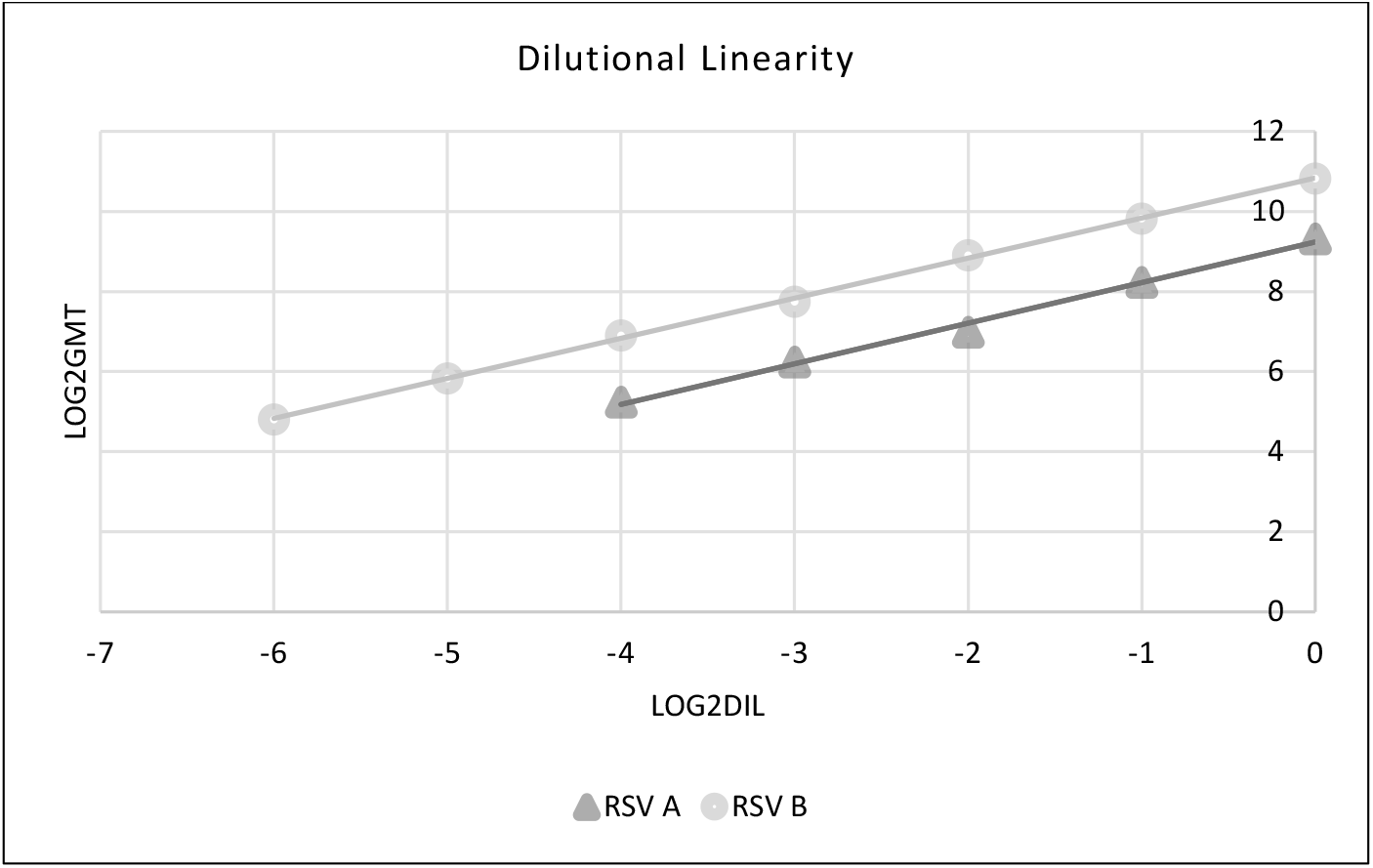
Dilutional linearity graph for RSV-A and B, showing the logarithm of the dilutions on the X axis and the logarithm of the obtained GMT on the Y-axis.

### Relative Accuracy

The assay is considered to have acceptable relative accuracy if the observed GMT of all replicates obtained for a sample is within ±50% to 200% of the expected titer. The MN assay is accurate from 1:1 to 1:16 dilution for RSV-A and from 1:1 to 1:64 dilution for RSV-B, as shown in **Table 1**.

**Table 1:**
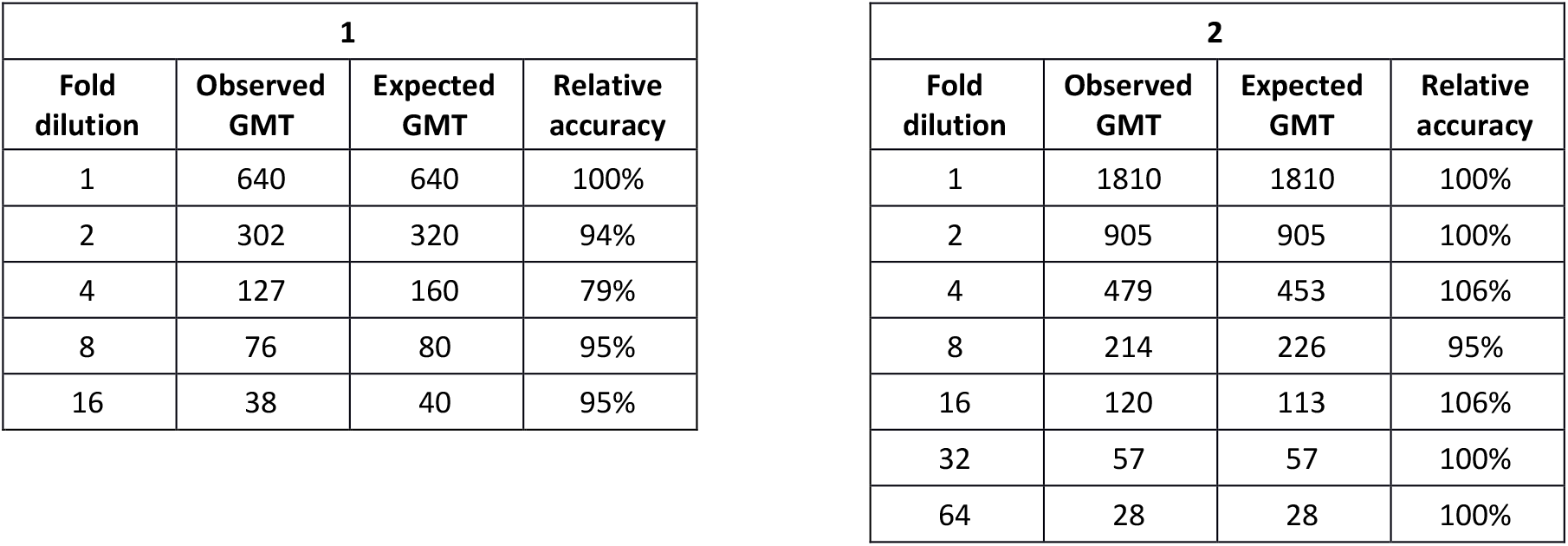
Relative accuracy results for RSV-A (1) and RSV-B (2).

### *Pre*cision

This parameter is usually indicated as the variance, standard deviation or coefficient of variation of a series of measurements (28).

### Repeatability

The assay is considered to have acceptable repeatability if all samples display a % geometric coefficient of variation (GCV) ≤65.5%. The results shown in **Table 2** indicate that the MN assay is repeatable from 1:1 to 1:16 dilution for RSV-A and from 1:1 to 1:64 dilution for RSV-B.

**Table 2:**
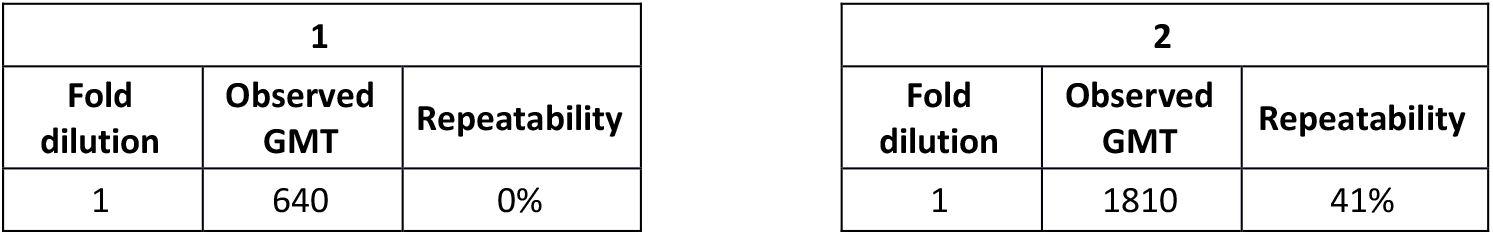
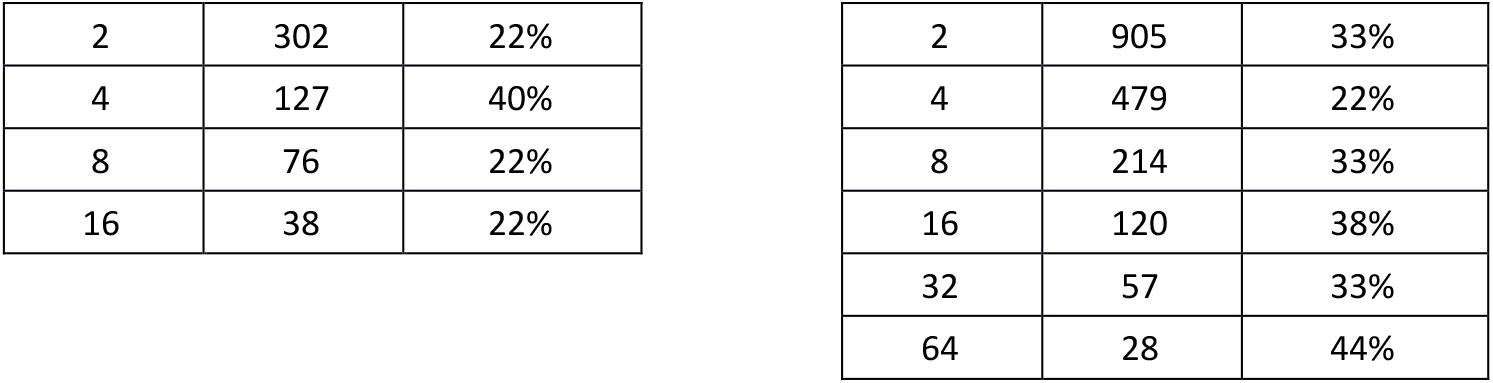
Repeatability (intra-run) results for RSV-A (1) and RSV-B (2).

### Intermediate Precision

The assay is deemed to be precise if all estimates of intermediate precision display %GCV ≤ 129.0%. The results reported in **Table 3** indicate that the MN assay is precise from 1:1 to 1:16 dilution for RSV-A and from 1:1 to 1:64 dilution for RSV-B.

**Table 3:** Intermediate precision results for RSV-A (1) and RSV-B (2).

| 1 |  |  |
| --- | --- | --- |
| Fold dilution | Observed GMT | Intermediate precision |
| 1 | 640 | 0.0% |
| 2 | 302 | 22.2% |
| 4 | 127 | 41.4% |
| 8 | 76 | 22.2% |
| 16 | 38 | 22.2% |

| 2 |  |  |
| --- | --- | --- |
| Fold dilution | Observed GMT | Intermediate precision |
| 1 | 1810 | 44.1% |
| 2 | 905 | 51.7% |
| 4 | 479 | 56.4% |
| 8 | 214 | 46.7% |
| 16 | 120 | 46.7% |
| 32 | 57 | 51.7% |
| 64 | 28 | 43.6% |

### Format Variability

The MN assay is considered to have acceptable format variability if all samples display %GCV ≤81.5%; the criterion was met for both RSV subtypes, as reported in **Table 4**.

**Table 4:** Format variability results for RSV A (1) and RSV B (2).

| 1 |  |  |
| --- | --- | --- |
| Fold dilution | Observed GMT | Intermediate precision |
| 1 | 640 | 0.0% |
| 2 | 302 | 15.2% |
| 4 | 127 | 27.8% |
| 8 | 76 | 15.2% |
| 16 | 38 | 15.2% |

| 2 |  |  |
| --- | --- | --- |
| Fold dilution | Observed GMT | Intermediate precision |
| 1 | 1810 | 29.5% |
| 2 | 905 | 34.2% |
| 4 | 479 | 37.2% |
| 8 | 214 | 31.1% |
| 16 | 120 | 31.1% |
| 32 | 57 | 34.2% |
| 64 | 28 | 29.2% |

### Range

For RSV-A, the LLOQ is set at 20 and the ULOQ is set at 640. For RSV-B, the LLOQ is set at 20, while the ULOQ is set at 2560.

### Specificity

Specificity is the ability of the assay to detect and distinguish the analyte of interest (38). To determine the specificity, positive samples for homologous (HS) and heterologous (HET) viruses were tested as reported in **Table S1**. The positive homologous sample showed a GMT with at least a four-fold difference from the heterologous samples. The negative sample showed negative titers in all measurements of this parameter. The results reported in **Table 5** indicate that the MN assay is specific.

**Table 5:** Specificity results for RSV-A and RSV-B showing the fold-change.

|  | HS/HET 1<br>(GMT) | HS/HET 2<br>(GMT) | HS/HET 3<br>(GMT) |
| --- | --- | --- | --- |
| RSV A | 32 | 32 | 32 |
| RSV B | 16 | 16 | 16 |

### Robustness

Two different cell suspension concentrations were used for each subtype: the standard concentration (3.0 × 10^4^ cell/mL) and a non-standard concentration (1.5 × 10^4^ cell/mL). Samples were tested in four replicates/plate, each by two different operators for each condition, thus obtaining sixteen titer values. The results obtained in each condition were aggregated to obtain eight RP values for each sample.

Plates were incubated for 60 min with the virus solution (standard condition) before the addition of the cell suspension; to assess the influence of the incubation time, plates were incubated with the viral solution for 45- and 75-min. Samples were tested in four replicates/plate by two different operators for each condition, thus obtaining twenty-four titer values. Results obtained from each condition were combined to obtain twelve RP values for each sample.

Regarding the cell suspension concentration, the %GCV was 19.65% for RSV-A and 17.40% for RSV-B, thus meeting the acceptance criterion set at ≤45%. With regard to the effect of the incubation time, the %GCV was 18.61% for RSV-A and 16.97% for RSV-B, thus meeting the acceptance criterion set at ≤45%. The above-mentioned results, together with the negative control, which showed negative titers, indicate that the MN assay is robust (**Table 6**).

**Table 6:** Robustness results for RSV-A and RSV-B, with standard deviation (SD) and % geometric coefficient of variation (GCV).

|  | Cell suspension concentration |  | Virus-serum mixture incubation time |  |
| --- | --- | --- | --- | --- |
|  | SD | %GCV | SD | %GCV |
| RSV A | 0.2 | 19.65 | 0.2 | 18.61 |
| RSV B | 0.2 | 17.40 | 0.2 | 16.97 |

## DISCUSSION

This study aims to provide guidelines and criteria for the establishment of a standard operating procedure for a high-throughput RSV neutralization assay. In the serological field, it is essential to have specific serological assays that can assess the efficacy of vaccines and antiviral drugs, evaluate monoclonal antibodies (mAbs) and establish new correlates of protections (CoP). The first step towards achieving this is to establish a common assay protocol (8). The definition of well-established CoP, especially mechanistic CoP, is a fundamental step towards designing better and more effective vaccines. Moreover, it is essential to understanding basic immunology and determining the susceptibility of a single individual or a population.

RSV has a high impact in infants, immunocompromised subjects and the elderly worldwide, in terms of death, hospitalization and the need for intensive care. Nevertheless, no FDA/EMA-approved vaccines against this disease have been licensed so far (39). This is probably due to the former reluctance of manufacturers to invest in new candidate vaccines, because of the high costs and the risk of serious adverse effects, such as the dramatic effects elicited by the first formalin-inactivated RSV vaccine in the 1960s. Currently, however, several vaccine candidates are in pre-clinical and clinical development. These include nasal live attenuated and temperature-sensitive vaccines, viral vector vaccines, and subunit or peptide vaccines (40).

The RSV MN assay is the most widely used method, since neutralizing antibodies play a key role in protecting against RSV infection and constitute the best correlates of protection. Thus, investigating the activity of nAb in depth could be of great value to the development of vaccines and new drugs against RSV, as happened in the case of SARS-CoV-2 vaccines immediately after the outbreak (41). The majority of vaccine candidates focus on the F protein in the pre-fusion conformation, which is known to be the most immunogenic form and elicits highly neutralizing antibodies. Binding assays, such as ELISA or Multiplexing assays in general, could be a valid surrogate for the neutralizing assays, as they are able to increase sample throughput. The main drawback is that they require the correct protein conformation of the F antigen (Pre-F). At present, the technologies for stabilizing and producing this type of antigen seem to be restricted, owing to patent pending issues. In addition, the establishment of a possible CoP for RSV should be based on the neutralizing response, since a high correlation between virus-specific functional neutralizing antibody titers and protection against the disease has been demonstrated. Thus, our aim was to describe the development of a validated assay and to underline the importance of a bioanalytical serological method able to specifically quantitate the presence of nAb after natural infection and/or vaccination.

The standardization and validation processes, performed according to well-defined international guidelines (28), are a key step to providing reliable results, especially when the assay is used to analyze samples from official clinical trials. In addition, it would be beneficial to use official WHO international standards, when available, in order to be able to properly standardize the assay and compare results from different laboratories and different assay formats, e.g. plaque reduction neutralization tests (IC50, IC80 or IC90) and ELISA-based neutralization tests. From our point of view, several crucial steps in the present method need to be highlighted and optimized in order to provide reliable results. Since this is a cell-based assay, the production and, more importantly, the stabilization of the live virus by means of a dedicated cryo-preservative, play a pivotal role in terms of the reproducibility of results. In addition, the cell line used for viral production and sample testing should be carefully evaluated, since susceptibility to infection can drive the assay in terms of high or low sensitivity. From our results, given the high homologies between RSV-A and B, the detection antibodies (against the F and N proteins) can be used for the read-out of both strains. However, the precise concentration for RSV-A and B independently remains to be investigated thoroughly. Finally, another aspect to examine is the infective dose, in terms of viral load, to be used in the MN assay. Since the rate of RSV infection in humans is quite high, and almost all people are found to be positive for the presence of nAb, it makes no sense to use a low infective dose, as was done in the case of SARS-CoV-2; indeed, at the beginning of the Covid pandemic, there was also an interest in distinguishing between responses in symptomatic and asymptomatic patients (42). The use of 100 TCID50/well has been acknowledged to be the right infective dose, even though higher infective doses may be used in the assay (200 or 300 TCID50/well). To conclude, the method of viral growth, stabilization and titration, as well as the RSV MN assay presented in this study, proves suitable for the quantification of the nAb titer in human serum samples, and may be used to assess the efficacy of new RSV candidate vaccines as well as in sero-epidemiological studies for the definition of CoP.

## AUTHOR CONTRIBUTIONS

CB and AM designed the experimental set-up. CMT and CB conceived the study, searched the literature and prepared the manuscript. IB, SL and SLE performed the neutralization assay. LB prepared and cultured the cell lines. ML was responsible for viral propagation and harvesting. CS and PP elaborated the results and performed statistical analyses. SM and CB performed statistical interpretation of results. EM and AM supervised the study.

