## Supplemental material for "Establishment and validation of a High-throughput Micro-Neutralization assay for Respiratory Syncytial Virus (subtypes A and B)"

**Table S1:** Serum samples and controls.

| | |
|---|---|
| **Positive control** | Antiserum to Respiratory Syncytial Virus WHO 1st International Standard (NIBSC) (HS) |
| **Negative control** | Negative human serum, Minus IgA/IgM/IgG |
| **Pool of homologous sera** | Normal serum, Discovery Life Sciences (cod. 150335) |
| **Heterologous sera** | Influenza Anti-A/Michigan/45/2015 (H1N1) (NIBSC) product code 17/106 (HET 1) |
| | Influenza Anti-A/Hong Kong/4801/2014 (H3N2) (NIBSC) product code 16/182 (HET 2) |
| | Influenza Anti-B/Brisbane/60/2008 (B/Victoria) (NIBSC) product code 16/192 (HET 3) |

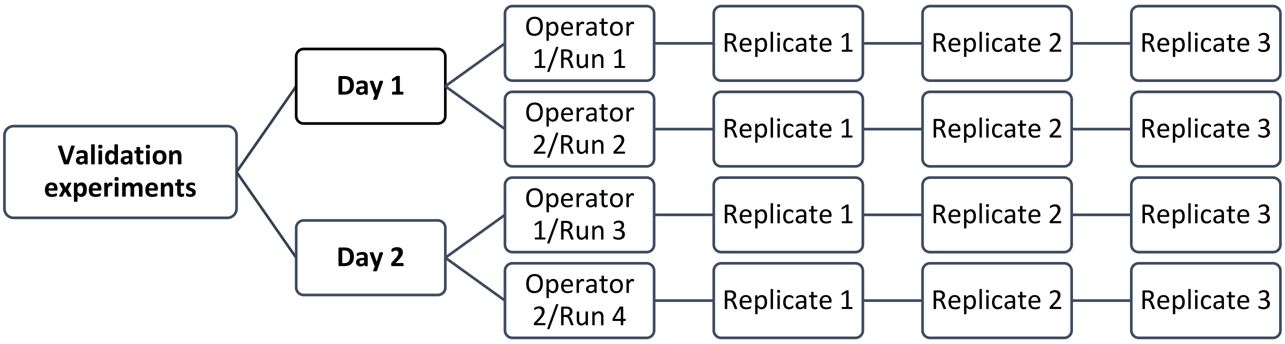


**Figure S1:** Validation Testing Design.
